# A Brain-Wide Genome-Wide Association Study of Candidate Quantitative Trait Loci Associated with Structural and Functional Phenotypes of Pain Sensitivity

**DOI:** 10.1101/2022.11.29.518322

**Authors:** Li Zhang, Yiwen Pan, Gan Huang, Zhen Liang, Linling Li, Zhiguo Zhang

**Author notes:** **Equal Contributions**, Dr. Li Zhang and Yiwen Pan. **Corresponding Authors**, Dr. Zhiguo Zhang.

## Abstract

Individual pain sensitivity is influenced by many factors, such as the differentiation of brain regional features and genetic variations. However, its heritability remains largely unclear. The present study conducted a brain-wide genome-wide association study (GWAS) to explore the genetic bases of structural and functional neuroimage phenotypes of pain sensitivity. In total 432 normal participants were firstly divided into high and low pain sensitivity groups according to laser quantitative test threshold and related brain regions’ gray matter density (GMD) features were identified. Consequently, GWAS was performed on each GMD phenotype using quality-controlled genotypes. Based on the heatmap and hierarchical clustering results from GWAS, the right insula was selected for further refined analysis in terms of sub-regions GMD and resting-state functional connectivity (rs-FC) phenotypes. The results indicate that the right insula GMD in the high sensitivity group is significantly lower than the low sensitivity group. Also, the TT/TC group at locus rs187974 may lead to a lower GMD in the right insula than the CC group. Meanwhile, loci at gene CYP2D6 may lead to a variation of rs-FC between the right insula and left putamen. In conclusion, our study suggests multiple candidate loci associated with pain sensitivity that may influence brain phenotypes.

## 1. Introduction

Pain is highly variable across individuals, and it is determined by many factors, such as the differentiation of regional features of the brain, genetic variation among the population, different psychiatric conditions, and even personalities (Rainville 2002; Coghill 2010; Zhang et al. 2021). Among these factors, the brain’s structure and function as well as genetic variation have been well-documented to have large effects on pain sensitivity. On one hand, the variation of brain structural and functional features may alter the way of pain perception since the brain processes the pain sensation. On the other, many single nucleotide polymorphisms (SNPs) have been found to have substantial impacts on pain sensitivity (Foulkes and Wood 2008). Therefore, to better understand individual differences in pain sensitivity, it is important to study the underlying brain imaging and genetic basis of pain sensitivity and to further provide mechanistic explanations of these differences.

Many brain imaging studies have shown that pain perception is largely dependent on individuals’ sensitivity to pain, and differences in pain sensitivity are often manifested by corresponding differences in the brain (Nielsen et al. 2009; Hu and Iannetti. 2016). Based on various types of brain imaging techniques, such as electroencephalography (EEG) and magnetic resonance imaging (MRI), former studies have successfully linked individual pain sensitivity to the variability in brain structure and function. Structural MRI studies have indicated that morphologic features of the brain, such as gray matter (GM) density and cortical thickness, are associated with pain sensitivity in healthy subjects (Erpelding et al. 2012; Emerson et al. 2014; Zhang et al. 2021). Further, some functional MRI (fMRI) studies indicated that regional blood oxygen level-dependent (BOLD) signal variability is inversely correlated with individual pain sensitivity within the ascending nociceptive pathway (primary and secondary somatosensory cortices), default mode network, and salience network (Rogachov et al. 2016). In addition, other research had established the heritable nature of both chronic pain and experimentally induced pain (Kato et al. 2006). In the meantime, several pain studies have shown an association between individual genetic variations and variability in pain perception. For example, twin studies have suggested that the heritability of different pain traits is about 50%, which implies that the variability of pain sensitivity is likely related to some specific genes (MacGregor et al. 2004). Though in recent years, many genes have been proposed to have associations with different types of pain, most studies conducted the analysis based on small groups of candidate genes (Young et al. 2012). Since pain perception is processed in the brain (Zou et al. 2021), it is of significance to reveal the genetic and neuroimage mechanism of pain sensitivity as well as to investigate their underlying relationships, such as grey matter density (GMD) and function connectivity (FC) from different pain sensitivity groups (Glahn et al. 2007; Ziv et al. 2010). However, a relatively large-scale imaging genetics database on pain sensitivity is still lacking. Consequently, further studies that unite brain features and genetic data on identifying pain sensitivity-related brain regions and genes are still very limited.

Imaging genetics for pain and other respects of brain-related research is an emerging interdisciplinary research field, which has shed light on how genetic variation influences brain structure and functions (Shen and Thompson 2019). One of the great advantages of using brain imaging genetics methods based on genome-wide association study (GWAS) is to combine brain imaging and genomics data with other phenotype data so as to reveal risk loci and genes that may be related to diseases and behavior traits (Visscher et al. 2017). GWAS makes use of high-throughput genotyping technology to relate a large number of SNPs to brain imaging quantitative traits (QTs) (Uffelmann et al. 2021), which can be more efficient than traditional case-control studies. However, only a few works were conducted via imaging genetics ways (Fontanillas et al. 2022), which impeded the exploration of neural mechanisms associated with genetic variation by fusing genes with imaging information.

Therefore, in this work, we focus on imaging genetics study using a self-collected dataset and mainly address the following three issues on pain sensitivity. First, identify brain regions that are significantly associated with laser pain sensitivity. Second, identify loci and genes that are related to the imaging QTs from brain structural and functional modalities. Thirdly, reveal the associations and effects among the genes, QTs, and pain sensitivity. Specifically, inspired by previous research (Shen et al. 2010), we present a brain-wide genome-wide association study using the laser quantitative test threshold to explore the brain and genetic architecture of pain sensitivity. GWAS was carried out on 432 normal participants, which were divided into high and low pain sensitivity groups according to their laser pain threshold from the laser quantitative test. Accordingly, brain structural and resting-state fMRI (rs-fMRI) images were acquired from an MRI scanner. A separate GWAS analysis using PLINK was completed for each of these phenotypes. Afterward, a heatmap and hierarchical clustering were used to demonstrate the significant associations between SNPs and phenotypes at threshold *p* < 10^-6^. Subsequent detailed analysis showed that several brain regions are associated with potential key loci from heatmap results. Since many previous studies reported that the insula plays a key role in pain perception (Ostrowsky et al. 2002), such a region and its subregions were selected for further analysis (Deen et al. 2011). A series of refined analyses using two-way analysis of variance (ANOVA) were performed to analyze the relationship between key loci, brain phenotype (GMD and FC), and pain sensitivity. Our results suggest that the brain-wide genome-wide association approach can reveal novel candidate genes and loci that may be related to pain sensitivity. The results show that individuals with a potential risk allele and high pain sensitivity at a key locus may have different structural and functional features in the corresponding brain regions. Overall, our work can help to understand the mechanism and physiology of pain sensitivity more comprehensively from the perspective of both imaging features and genetics.

## 2. Materials and Methods

### 2.1 Participants

We recruited 432 healthy participants (259 females, age: 21.30 ± 3.99 years) via college and community advertisements for the experiment. Twenty left-handed participants were excluded due to differences in brain morphology and function among left-handed and right-handed participants (Pool et al. 2015; Margiotoudi et al. 2019; Wiberg et al. 2019). Before the experiments, participants were carefully screened to ensure that they had no history of chronic pain, neurological diseases, cerebrovascular diseases, coronary heart disease, or mental disorders, and they had no contraindications to MRI examination. The study was proved by the local ethics committee and all participants gave their written informed consent before participating in the study.

### 2.2 Extraction and genotyping

Five-milliliter blood samples were collected from participants for DNA extraction. Genotypes of SNPs were determined with the CapitalBio Technology Corporation Precision Medicine Research Array based on the Affymetrix platform. After the above processing, 748,659 SNP markers were collected as genome-wide genotype information.

### 2.3 Pain sensitivity definition

The pain sensitivity of all the participants was measured as laser pain thresholds using quantitative sensory testing before the MRI scan. A series of infrared neodymium yttrium aluminum perovskite (Nd: YAP) laser stimuli were delivered to the back area between the thumb and index finger of a participant’s left hand. The laser pain threshold was acquired using quantitative sensory testing. The measurement was started from an energy level of 1 J with a 0.25 J increase at each stimulus. After each stimulus, a participant was asked to report the pain rating from 0 (no pain) to 10 (unbearable pain). When a rating of 4 was reported, the corresponding energy level was recorded as the laser pain threshold of the participant. For each participant, the laser pain threshold was averaged from two independent measurements conducted in one hour.

The values of the laser pain threshold were normalized by z-score normalization to determine the pain sensitivity threshold to meet the normal distribution. Based on the above threshold, the participants were first sorted from the low pain threshold (high sensitivity to pain) to the high pain threshold (low sensitivity to pain) and then divided into two groups based on the zero threshold.

### 2.4 MRI data acquisition

MRI data were obtained on a 3.0-Telsa system (GE Medical Systems, Milwaukee, WI, USA) at the Brain and Cognitive Neuroscience Research Center, Liaoning Normal University, Dalian, China. A standard birdcage with an 8-channel head coil, along with restraining foam pads, was used to minimize head motion and scanner noise. High-resolution structural T1-weighted images were collected using a three-dimensional magnetization-prepared rapid gradient echo (3D-MPRAGE) sequence with the following imaging parameters: flip angle = 8°; field of view = 256 × 256 mm^2^; data matrix = 256 × 256; in-plane resolution = 1 × 1 mm^2^; slices = 176; slice thickness = 1 mm. Ten-minute resting-state functional images were acquired using an echo-planar-imaging sequence with the following imaging parameters: repetition time = 2000 ms; echo time = 29 ms; flip angle = 90°; field of view = 192 × 192 mm^2^; data matrix = 64 × 64; in-plane resolution = 3 × 3 mm^2^; slice thickness = 3 mm. During the resting-state fMRI data acquisition, participants have presented with a screen with a white crosshair in the center of the black background. They were instructed to relax, remain awake, and not think about anything with their eyes open.

### 2.5 Processing of structural MRI data

The raw DICOM scans were converted into the Neuroimaging Informatics Technology Initiative format using MRICRON software after review. We performed the preprocessing steps using CAT12 toolboxes under SPM12 (Statistical Parametric Mapping, http://www.fil.ion.ucl.ac.uk/spm) with the default setting. Voxel-based morphometry (VBM) (Good et al. 2001) was used to process and extract brain-wide target MRI imaging phenotypes. The structural images were segmented into GM, white matter (WM), and cerebrospinal fluid (CSF) components by applying a registration to the Montreal Neurological Institute (MNI) stereotactic space and subsequent non-linear deformation. The non-linear deformation parameters were calculated via the inbuilt high dimensional Diffeomorphic Anatomical Registration Through Exponentiated Lie (DARTEL) Algebra algorithm (Ashburner 2007). Then, the warping functions generated by DARTEL were used to spatially normalize the GM segments and modulate them by the Jacobian determinant. Finally, the normalized GM images were smoothed using an 8-mm full width at half maximum Gaussian kernel. After the above processing, the total GM volume (GMV), WM volume, and CSF volume were obtained from the smoothed images in the CAT12 toolboxes and saved in the total intracranial volume (TIV).

A regression analysis was performed using a general linear model to examine the relationship between GMD across the whole brain and pain sensitivity (Emerson et al. 2014). Age, gender, and TIV were regressed out as covariates. Then brain regions with a significant relationship between GMD and normalized pain threshold (*p* < 0.05) were extracted as relevant regions of interest (ROIs) for subsequent association analysis.

### 2.6 Processing of functional MRI data

The rs-fMRI data were preprocessed with Data Processing & Analysis for Brain Imaging (DPABI) (http://www.rfmri.org/dpabi) (Yan et al. 2016). First, the initial 10 volumes of the time series were removed due to the stability of the gradient magnetic field and participants’ adaptation to the scanner circumstances. The remaining images were slice-timing corrected, realigned, and resliced to correct for head motion. Participants were excluded if their head motion was more than 2.0 mm of translation or 2.0° of rotation throughout scanning for further analysis (sixteen participants were excluded). Then all fMRI images were spatially normalized into the MNI space using the echo-planar imaging (EPI) template, and each voxel was resampled to isotropic 3×3×3 mm^3^. After normalization, all data sets were smoothed with a Gaussian kernel of 8×8×8 FWHM. Next, the following nuisance covariates from each voxel’s time course were regressed out, including 24 head motion parameters (Friston et al. 1996), WM, and CSF signals, as well as the linear trend. Subsequent data required temporal filtering in a frequency range of 0.01 to 0.1 Hz to reduce the effects of low-frequency drift and high-frequency noise.

### 2.7 Brain parcellation and extraction of multi-modality imaging phenotypes

The automated anatomical labeling (AAL) atlas 3 (Rolls et al. 2020) was used to create the ROI level measurements. For structural MRI data, the GMD at the level of 170 ROIs was extracted from GMD images as structural image phenotypes. For resting-state MRI data, time series of whole-brain were obtained for resting-state functional connectivity (rs-FC) phenotypes. More specifically, for each participant, the time series was extracted as the mean value across all voxels within that ROI. For each ROI, the Pearson correlated coefficient (r-value) between itself and all other ROIs in the AAL3 template was calculated by the voxel-voxel method (Allen et al. 2014). Then, we converted the r-values to z-values by applying Fisher’s r-to-z transformation (Liu et al. 2019). Finally, the z-values between pairs of ROI were used as the rs-FC phenotypes.

### 2.8 Quality control

To handle possible defects of genetic data, the following quality control (QC) steps were performed on these genotype data using the PLINK software package (v1.9) (https://zzz.bwh.harvard.edu/plink/) (Purcell et al. 2007). Firstly, only SNPs located at autosomes were included in this study and 27,932 SNPs at sex chromosomes were removed. Secondly, SNPs were excluded if they could not meet any of the following criteria: (1) call rate per SNP ≥90% (46,803 SNPs removed), (2) minor allele frequency (MAF) ≥5% (319,751 SNPs removed), and (3) Hardy-Weinberg equilibrium test of p≤10^-6^ (752 SNPs removed). Participants were excluded from the analysis if any of the following criteria was not satisfied: (1) call rate per participant ≥90%; (2) gender check (5 participants were excluded). Population stratification analysis (Freedman et al. 2004) was performed by applying the principal component analysis (PCA), which is considered to be a very useful method of adjusting spurious associations. This method enables us to obtain the ethnic information of the sample and to determine possible ethnic outliers. After the QC procedure, 407 out of 412 participants and 353,421 out of 748,659 SNP markers remained in the analysis.

### 2.9 Genome-wide association analysis

The GWAS analysis was conducted separately on structural and rs-FC phenotypes after the QC procedures of the SNP data. Before that, the phenotypes were adjusted for the baseline age, gender, TIV or head motion, and PCA parameters. We applied the PLINK software package to calculate the main effects of all SNPs on the corresponding quantitative imaging phenotype with the QT association option. An additive SNP effect was assumed, and the empirical p-values were based on the Wald statistics (Purcell et al. 2007).

To investigate the GWAS results and data reduction for subsequent refined analyses, we employed a heatmap to visualize the association pairs between identified SNPs and corresponding imaging phenotypes at different statistical significance levels (Sloan et al. 2010). A color map was used to represent various significance levels. Traditionally, the association between an SNP and an imaging phenotype can be regarded as significant if *p* < 10^-6^ is found (Shen et al. 2010). Based on the obtained heatmap result, a hierarchical clustering method was applied to identify the similarity among SNPs and imaging phenotypes, respectively. The clustering was completed using the Euclidean distance method to define the dissimilarity between two nodes. The distance average among all pairs of objects in two clusters was used to measure the similarity between the two clusters. In this way, two dendrograms can be obtained to show the clustering results for imaging genotypes and SNPs respectively. All association results surviving the significance threshold of *p* < 10^-6^ were saved and prepared for additional pattern analysis.

Following many previous pain-related studies which reported that the insula plays a key role in the emotional-affective dimension of pain and for pain modulation (Ostrowsky et al. 2002), the GMD of the right insula was selected for further refined analysis. It is because the number of significant SNP associated with the right insula was highest. A Manhattan plot and a quantile-quantile (Q-Q) plot were used to visualize the GWAS results for the right insula. Moreover, since former works show that each subregion of the insula has different cell structure and functional distributions in pain perception (Mazzola et al. 2012; Wiech et al. 2014; Lu et al. 2016), bilateral insulae were further divided into three subregions in subsequent analyses through data-driven clustering techniques (Ben et al. 2011). The above method was performed in both structural and resting-state functional image features. Based on (Menon et al. 2020), we obtained 3 insula subregions in the right hemisphere, including the posterior insula (PI), the dorsal anterior insula (dAI), and the ventral anterior insula (vAI), by merging the related regions for further statistical analysis.

### 2.10 Refined analysis

After extracting the genotype information, refined analyses were further conducted to quantitatively determine the difference of multi-modality phenotypes between various genotype groups. More specifically, the participants were divided into three groups based on different loci, including two homozygous groups and one heterozygous group. Due to the uneven distribution of the number of participants in different groups, the homozygous group with a small number of participants was combined with the heterozygous group as a risk allele-carrier group (Liu et al. 2019). After grouping participants by genotype, a two-sample t-test was used to determine the association between genotype and phenotype characteristics. Groups of high and low pain sensitivity were then divided, and a two-sample t-test was used to examine the association between pain sensitivity and phenotypic features. Finally, a two-way ANOVA (2×2 ANOVA: pain sensitivity × genotypes) was performed to examine the effect of target SNP and pain sensitivity on both GMD and rs-FC phenotype of target ROI. All statistical analyses were performed using SPSS 22.0 software.

## 3. Results

### 3.1 Sample characteristics after QC

After QC procedures of the genotyping data, 407 out of 432 recruited participants remained in the present study. Fig. 1 shows the distribution and the fitting result using a Gaussian distribution of the normalized pain threshold (after Fisher’s r-to-z transformation) of the 407 participants. The normalized pain thresholds are sorted from low (high pain sensitivity) to high (low pain sensitivity) with a roughly normal distribution. The figure demonstrates that the number of participants with high pain sensitivity is relatively greater than that with low pain sensitivity. Then structural and functional MRI data from all these 407 participants were preprocessed, and the preprocessed GMD and rs-FC values were used as phenotypes for subsequent GWAS analysis. Table 1 illustrate the demographic information of the sample analyzed for both structural and resting-state functional studies. Among the participants, the genders of the participants are significantly different (p = 0.027) between the high and low sensitivity groups after the independent-sample t-test when dealing with structural phenotypes. In the subsequent GWAS analyses, age, gender, TIV, or head motion were considered as the covariates.

**Fig. 1.**
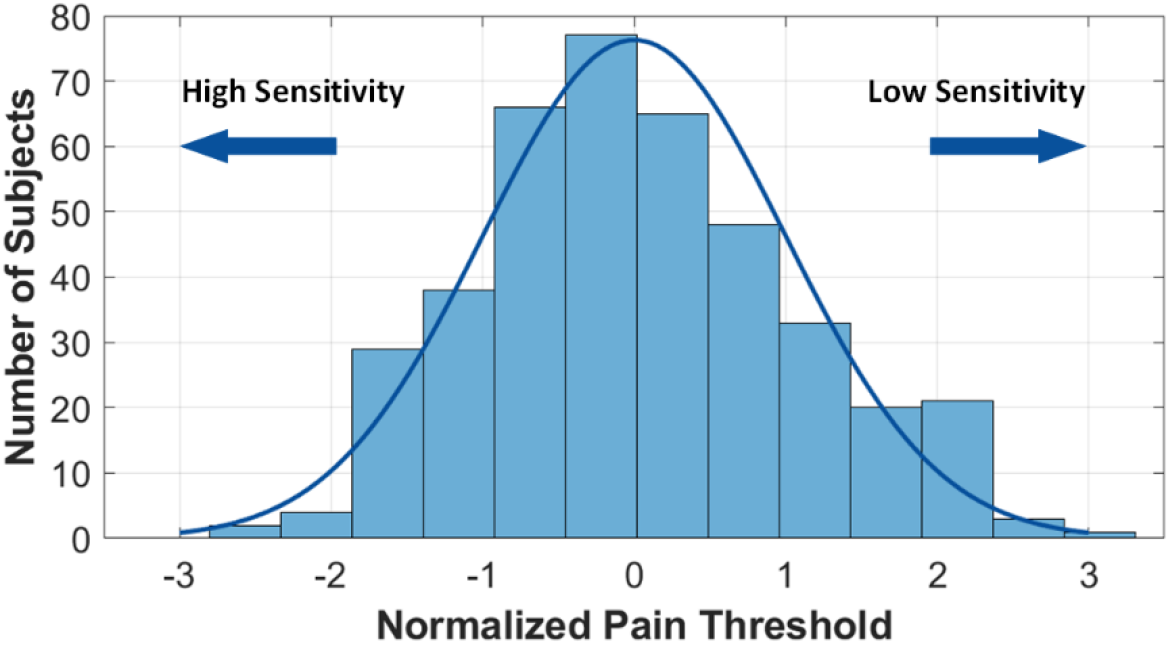
Histogram with a normal distribution fit of normalized pain threshold (after Fisher’s r-to-z transformation) of 407 participants. The blue curve is the fitting result of the data with a Gaussian distribution.

**Table 1.** Demographic information and the total number of participants are included in each analysis. After quality control, GMD was extracted as a structural phenotype from the images of 407 participants, and rs-FC values were extracted as a functional phenotype from the resting-state MRI of 390 participants. Basic demographic information is shown for both groups of subjects.

| Category | Structural phenotypes (407 participants) |  |  | rs-fMRI phenotypes (390 participants) |  |  |
| --- | --- | --- | --- | --- | --- | --- |
|  | High | Low | p-value | High | Low | p-value |
| Number of participants | 216 | 191 | - | 210 | 180 | - |
| Gender (M/F) <sup>a</sup> | 74/142 | 86/105 | **0.027\*** | 73/137 | 79/101 | 0.067 |
| Age (years, mean±SD) <sup>b</sup> | 21.1±3.9 | 21.4±4.1 | 0.508 | 21.2±3.9 | 21.2±3.6 | 0.869 |
| Education (year, mean±SD) <sup>b</sup> | 14.6±2.0 | 14.5±1.9 | 0.746 | 14.6±2.1 | 14.6±2.0 | 0.765 |

Note: <sup>a</sup> chi-square test; <sup>b</sup> two-sample t-test; \* p-value <0.05

### 3.2 Extraction of ROIs associated with pain

Multiple regression of normalized pain threshold and GMD in unrestricted ROIs in the whole brain was performed based on SPM12 (adding age, gender, and head motion as covariates). Fig. 2 shows that the activated brain regions were basically symmetrical. Moreover, both left and right insulae correlated with normalized pain threshold significantly (*p* < 0.05, uncorrected). Then GMD of 98 brain regions was extracted as structural phenotypes for subsequent GWAS analysis.

**Fig. 2.**
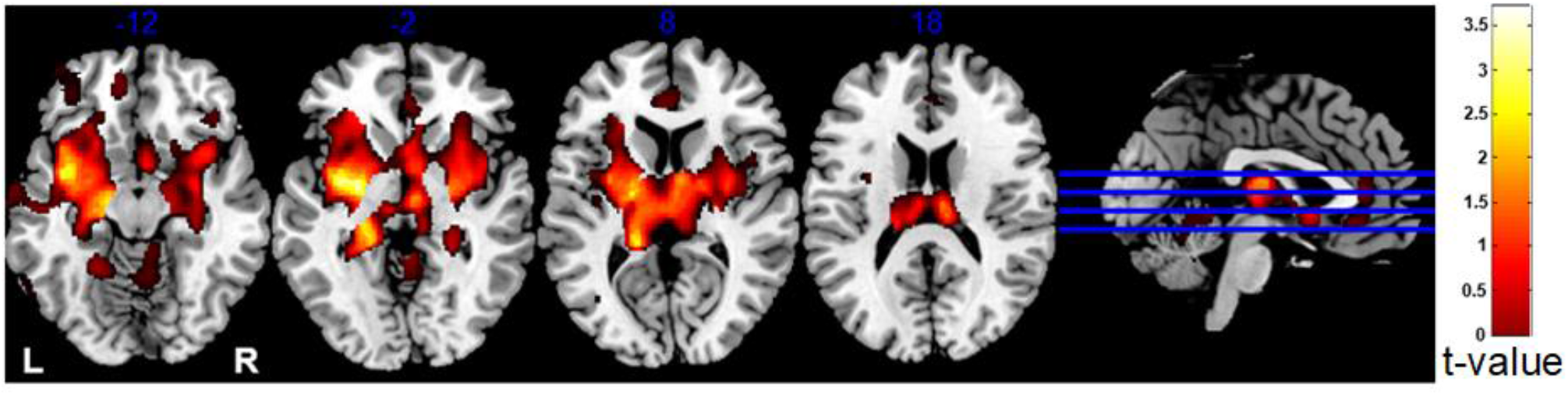
Relationship between normalized pain threshold and GMD after multiple regression. Normalized pain threshold is positively correlated with GMD, while pain sensitivity is inversely related to GMD.

### 3.3 GWAS of GMD

The heatmap shown in Fig. 3 illustrates the imaging genetics associations at a significance threshold of *p* < 10^-6^, which were discovered by GWAS analysis of 98 structural phenotypes, i.e., brain imaging QTs. The names of identified brain regions and SNPs are shown accordingly. On the heatmap, significant associations between imaging phenotypes and SNPs are marked with an “x” if *p* < 10^-6^. At this significance level, 50 strong SNP-QT associations (blocks labeled with “x”) were identified in the GWAS analyses, and 46 SNPs were involved in these associations. Two dendrograms for the hierarchical clustering results of imaging genotypes and SNPs can be obtained along with the x-axis and y-axis respectively. The color bar on the left side of the heatmap encodes the chromosomes for the corresponding SNPs.

**Fig. 3.**
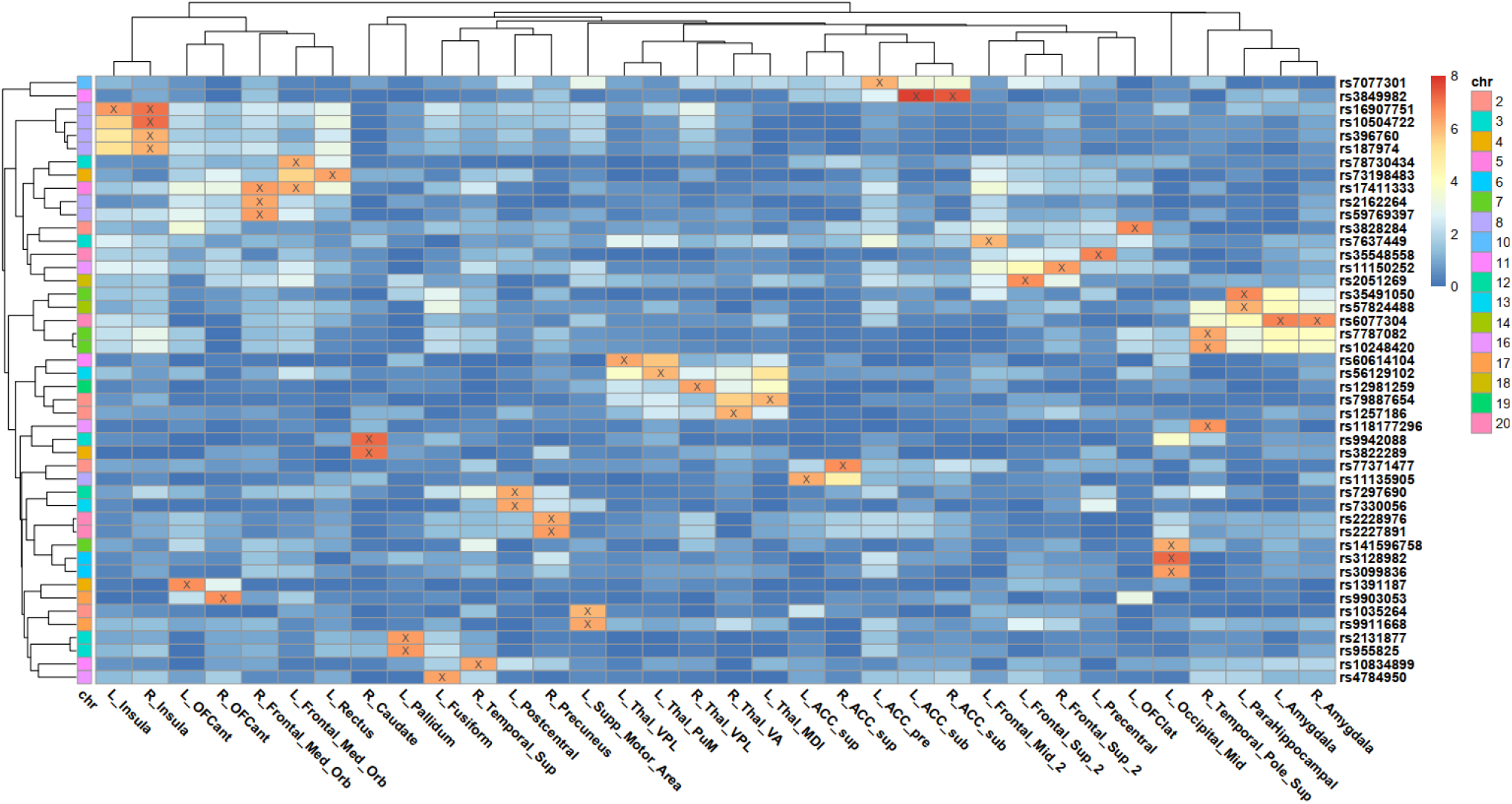
Heatmap of SNP associations with QTs at the p≤10^-6^ significance threshold. The blocks with “x” indicate the significance level of p≤10^-6^. Only significant SNPs and QTs are shown in the heatmap. Dendrograms from the x-axis and y-axis were derived from hierarchical clustering for QTs and SNPs respectively. The color bars on the right side encode the p-values and corresponding chromosomes for SNPs respectively.

From Fig. 3, we can see that several imaging phenotypes were identified to have strong associations with target SNPs in the GWAS analyses. From the imaging dimension, bilateral insulae and amygdala regions, which are considered to be related to pain perception and modulation can be identified. Also, it is interesting to find that a group of regions such as caudate, pallidum, and supplementary motor area can also be identified. It suggests that these regions are QTs associated with individual pain sensitivity. Observed from the genomic dimension, several SNPs can be identified from the results, including rs10504722, rs16907751, rs396760, and rs187974. It can be seen from the heatmap that all these four SNPs are associated with the right insula.

The hierarchical clustering results shown in Fig. 3 indicate that many pairs of left and right measures of the same region were clustered together as shown in the imaging dimension (x-axis). This suggests the symmetric relationship between these phenotypes and genetic variations. Moreover, several SNP groups from the genomic dimension (y-axis) in the same color can be identified from chromosome 8 (rs10504722, rs16907751, rs396760, and rs187974). This is probably caused by the linkage disequilibrium (LD) effect that increases the likelihood of non-random association of different alleles.

The right insula was selected as a target QT for further detailed analysis because it has the largest number of SNPs and a strong association with these SNPs. In Fig. 4(a), a Manhattan plot of the GWAS for the target QT is shown. The x-axis of the Manhattan plot denotes different chromosomes, and the y-axis denotes the p-value of all SNPs. The Manhattan plot indicates that several potential SNPs located at chromosome 8 may be associated with pain sensitivity. Correspondingly, the Q-Q plot of the right insula is shown in Fig. 4(b). From the figure, we can see that, for most of the p-values, the observed p-values from GWAS are almost the same as expected from the null hypothesis. The p-values in the upper tail of the distribution show a significant deviation. It suggests strong associations between these SNPs and the GMD of the right insula. Fig. 4(c) shows the relationship between pain sensitivity and GMD in the right insula, and we can see that participants with higher normalized pain thresholds were found to have larger GMD in the target QT relative to participants with higher intensity ratings. It means that in the right insula, GMD and pain sensitivity are negatively correlated. The insula was further divided into three subregions in the right hemisphere to identify the morphological abnormalities. The GMD of each insula subregion was extracted and used to correlate with genotypes separately.

**Fig. 4.**
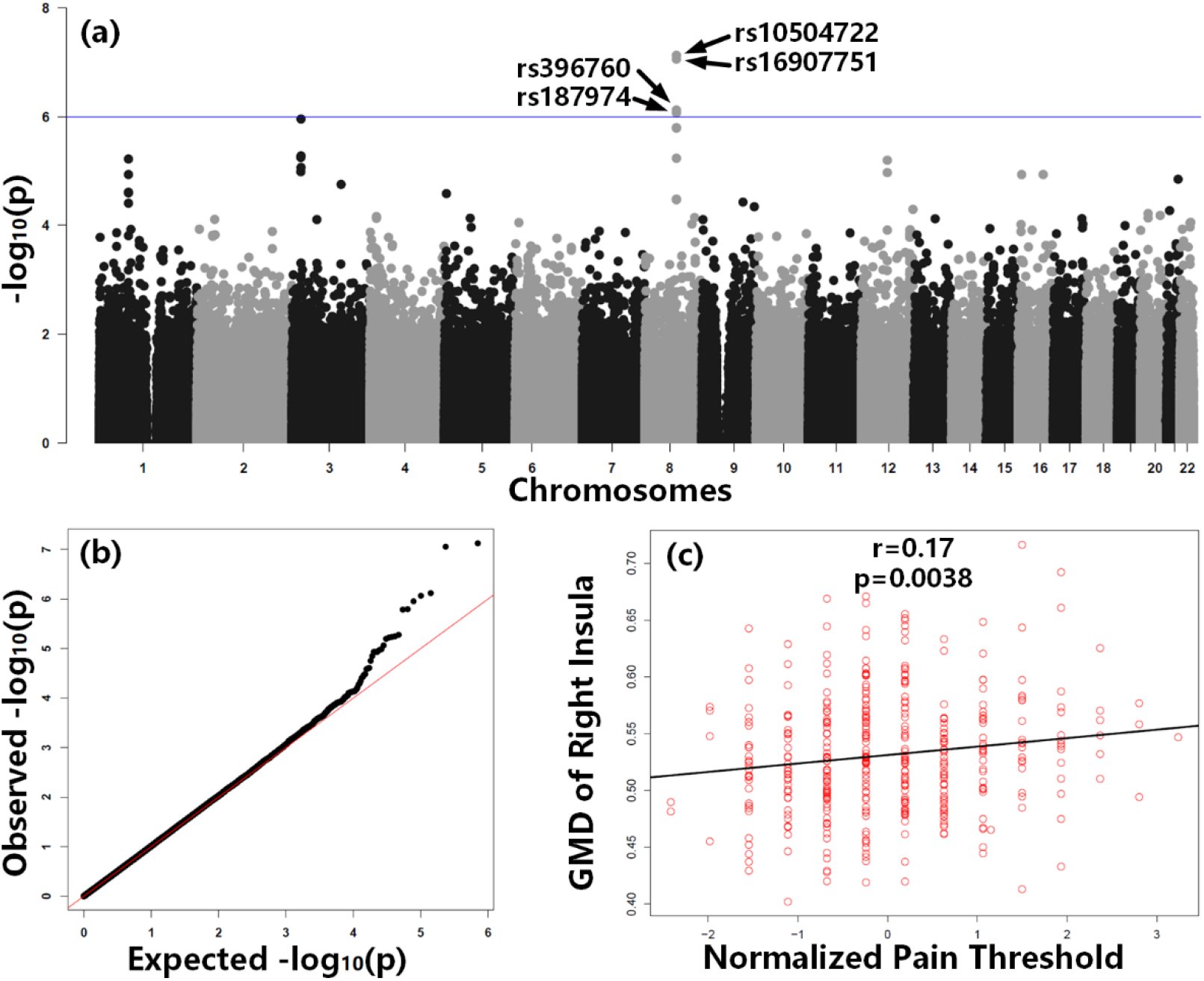
Manhattan and Q–Q plots of GWAS of QT as the GMD of right insula, and correlation between GMD and normalized pain threshold. (A) The blue horizontal line in the Manhattan plot indicates p = 10^-6^. Four significant SNPs are marked with their names. (B) The Q-Q plot of the right insula indicates the distribution of the observed p-values in this sample versus the expected p-values under the null hypothesis of no association. (C) Correlation analysis revealed a positive correlation between GMD and normalized pain threshold (r = 0.17, p = 0.0038).

### 3.4 Refined analysis for SNP with structural phenotype

A target SNP, rs187974, corresponding to the right insula is selected from the heatmap for further imaging analysis. Additionally, since the number of participants with a TT genotype in rs187974 is too small, participants with TT and TC genotypes were combined into the TX group. As shown in Fig. 5, a two-way ANOVA using SPSS was performed to analyze the effect of the pain sensitivity group and rs187974 genotype on GMD of the right insula (n=407; 202 high pain sensitivity (39 TX, 173 CC); 183 low pain sensitivity (36 TX, 147 CC); 2 missings). In the figure, the mean and standard deviation (SD) of GMD are shown. A main effect of genotype shows that the GMD of the CC group (as shown in light blue) is elevated more than that of the group of T-allele carriers (as shown in dark blue) of the right insula (*p* = 0.0105). Moreover, the participants with high pain sensitivity have lower GMD than those with low pain sensitivity (*p* = 0.0487). Accordingly, it may suggest that if a participant is high-pain sensitive and carries the potential high-risk allele T at rs187974, the GMD of the subject may be lower than others.

**Fig. 5.**
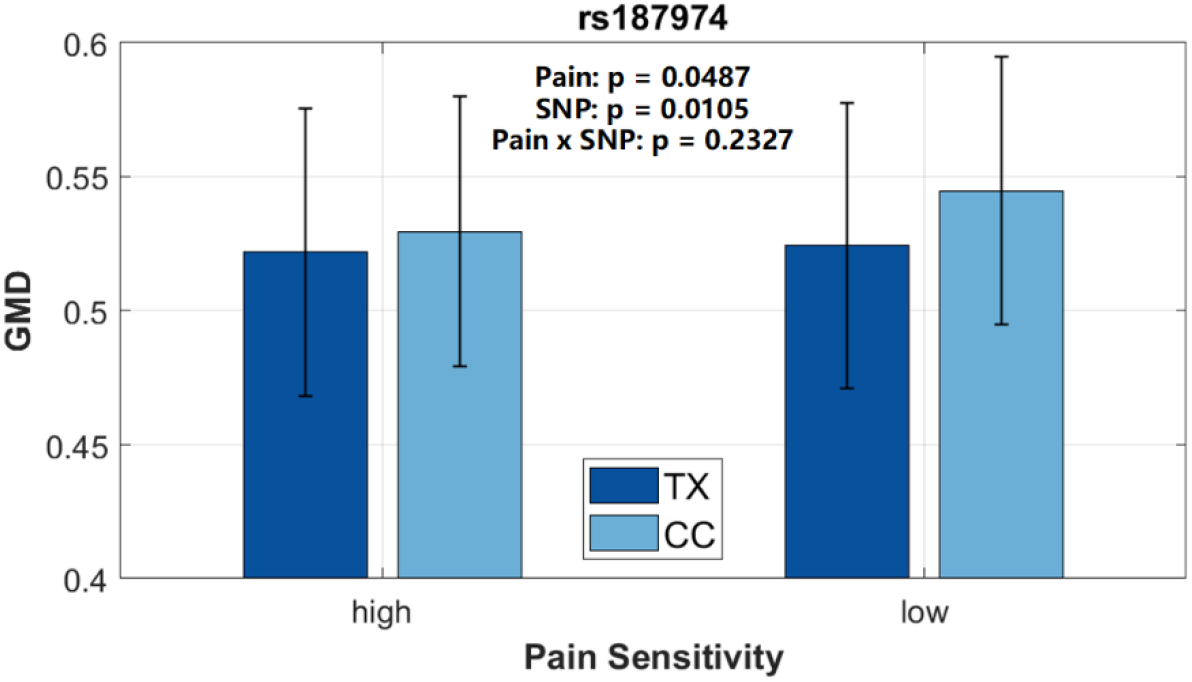
Refined analysis of GMD of the right insula in relation to rs187974 and pain sensitivity group. A two-way ANOVA was performed to examine the effect of rs187974 and pain sensitivity on GMD of the right insula. The bars and error bars indicate the mean and SD of GMD.

After performing the GWAS analysis on GMD of subregions of the right insula and SNP data, only four SNPs (rs10504722, rs16907751, rs396760, and rs187974) were found to be significantly associated with their corresponding dAI. These SNPs were consistent with those obtained when the entire insula was analyzed, suggesting that the main effect of GMD and SNP association is on the dAI. As shown in Fig. 6, the homozygous group with a small number of participants is also integrated with the heterozygous group. The results illustrate that the participants with high pain sensitivity carrying the T allele in the rs16907751 and rs187974 SNPs (*p* = 0.0002 and *p* = 0.0079 respectively) may have lower GMD in dAI than others. In addition, paired comparisons showed that the GMD of the dAI was lower in participants with risk allele A than in participants with C-allele, and the GMD of subjects with higher sensitivity was lower than those with lower pain sensitivity. This indicates that participants with high pain sensitivity carrying A-allele in rs396760 SNP (*p* = 0.0101) may have lower GMD in the dAI than others. Meanwhile, the result shows that participants with high pain sensitivity carrying risk C-allele in rs10504722 (*p* = 0.0037) have lower GMD of dAI than others.

**Fig. 6.**
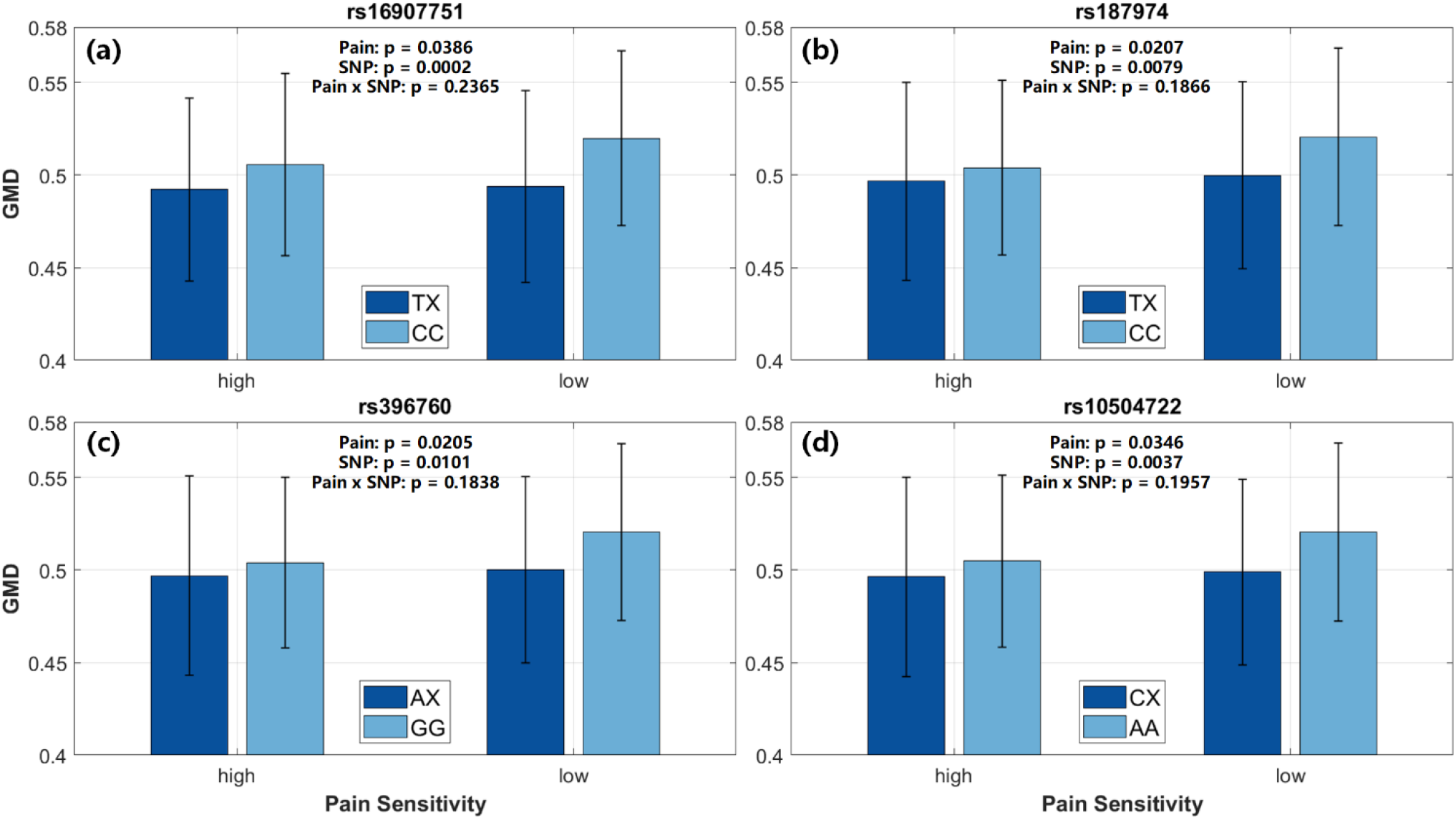
Refined analysis of GMD of subregions of right insula about corresponding SNPs and pain sensitivity group. Two-way ANOVAs were conducted to examine the effect of the four SNPs and pain sensitivity on GMD of the dorsal anterior insula (dAI). The main effect of the pain sensitivity group and (A) rs16907751; (B) rs187974; (C) rs396760 and (D) rs10504722. The p-values for the main effect of the pain sensitivity group and SNP are shown in the plot. The bars and error bars indicate the mean and SD of GMD.

### 3.5 Refined analysis for SNP with functional phenotype

The subjects’ rs-FC of pain-related brain regions are highly correlated with individual pain sensitivity (Icenhour et al. 2017; Monroe et al. 2018). Since the insula is a functionally diverse brain region that plays a central role in multiple aspects of pain processing (Segerdahl et al. 2015), it is crucial to investigate the rs-FC variations among subjects with different pain sensitivity and its genetic basis. Based on the previous structural phenotype analysis that focused on the right insula, we still used the right insula as the seed point to get the correlation coefficient as the rs-FC value between the right insula and other brain regions. After GWAS analysis, we found that the rs-FC between the left putamen and right insula can be identified as a significant QT that correlated to SPNs rs2267447 and rs1081003. These two SNPs were extracted for subsequent statistical analysis. It is worth noting that both SNPs belong to the same gene named CYP2D6 according to Single Nucleotide Polymorphism Database (dbSNP, https://www.ncbi.nlm.nih.gov/snp/).

We further conducted the statistical analysis using a two-way ANOVA to examine the effect of target SNPs and pain sensitivity groups on the rs-FC between the right insula and the left putamen. The results are shown in Fig. 7. Firstly, both Figs. 7 (a) and (b) indicate that participants with high pain sensitivity had significantly lower rs-FC values than low pain sensitivity for both target SNPs (*p* = 0.0163 and *p* = 0.0246). In Fig. 7(a), the main effect of the rs2267447 genotype across all participants was significant for the rs-FC value. More specifically, the rs-FC of the CC group was significantly lower than the TT group in the rs2267447 SNP (*p* = 6.17 × 10^-5^). In Fig. 7(b), we can also find that the rs-FC of the AA group was lower than the GG group in the rs1081003 SNP (p = 6.08 × 10^-5^). Both results suggest that participants with high pain sensitivity carrying C-allele of rs2267447 and A-allele rs1081003 may have a lower rs-FC value between the right insula and left putamen than rs-FC.

**Fig. 7.**
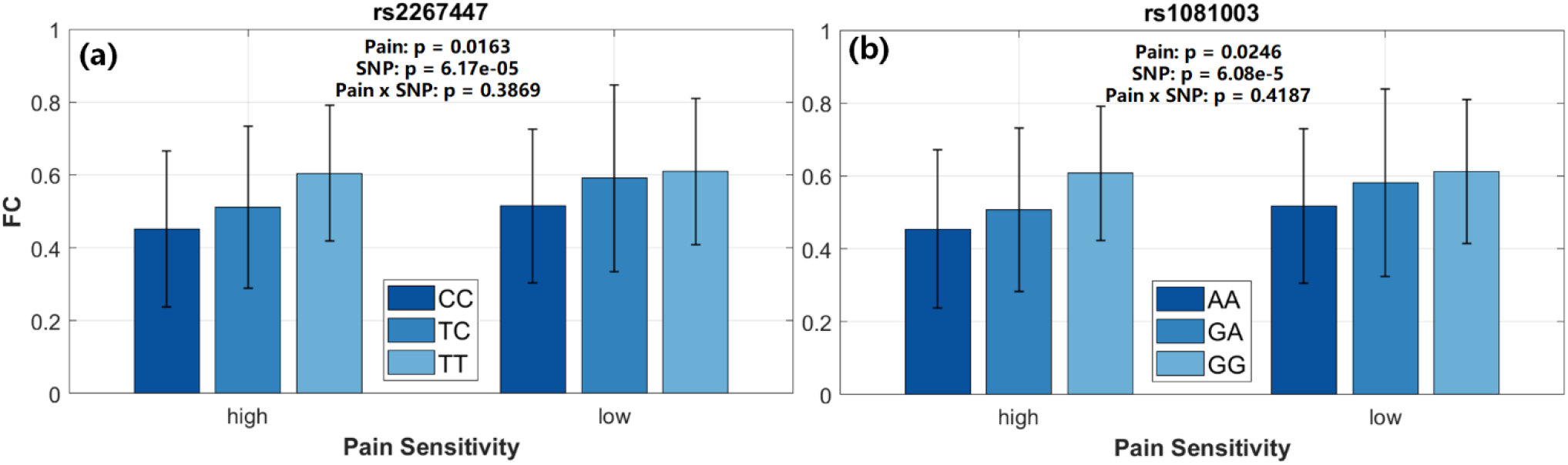
Refined analysis of the rs-fMRI phenotype of right insula about corresponding SNPs and pain sensitivity group. Two-way ANOVAs were applied to examine the effect of the two SNPs and pain sensitivity on rs-FC between the right insula and the left putamen. The main effect of the pain sensitivity group and (A) rs2267447 and (B) rs1081003. Both SNPs belong to the gene CYP2D6. The bars and error bars indicate the mean and SD of rs-FC.

## 4. Discussion

### 4.1 Imaging phenotype definition

Pain sensitivity is a characteristic that affects the perceiving way of painful stimuli for different subjects. Though enormous progress has been made from previous studies, the reason why different individuals have different levels of pain sensitivity, and what factors contribute to pain sensitivity individual differences are still open and challenging questions. Aiming these, we employed a whole-genome-whole-brain method in this paper to systematically identify candidate quantitative trait loci associated with brain imaging phenotypes of pain sensitivity. The data analysis method can be summarized into three major steps: (1) preprocessing and quality control; (2) GWAS of brain structural phenotypes, and (3) GWAS of brain rs-FC phenotypes. This approach is similar to other imaging genetics research (Shen et al. 2010; Stein et al. 2010).

To investigate the brain structural and functional basis of pain sensitivity, the values of GMD from 170 ROIs were extracted. From the results shown in Fig. 2, there are 98 brain regions can be identified to be related to subjects’ pain threshold. Most of these regions are located in the insula, basal ganglia, thalamus, and hippocampal regions. The strongest association area with pain threshold from our experiment is located in the insula cortex. These results are promising because a number of past studies have indicated that these regions play key roles in pain perception and modulation (Barker 1988; Chudler and Dong 1995; Starr et al. 2009; Borsook et al. 2010; Lu et al. 2016). For example, Lu has reported that the anterior portion of the insular cortex is critical for integrating and competing for multiple information to create awareness of the pain (Lu et al. 2016). Other studies have shown potential pathways via basal ganglia with respect to possible pain processing (Barker 1988; Chudler and Dong 1995). Along with the findings from our results, these brain regions could be potential therapeutic targets for effective and individualized pain modulation, such as precision anesthesia and pain relief.

### 4.2 GWAS of structural brain phenotypes

In Fig. 3, the heatmap and hierarchical clustering have been used to discover the SNP-by-ROI associations and the grouping results of different SNPs. One reason for using a heatmap is that it can give an intuitive representation of the associations between SNPs and target brain regions. These results also align well with the associated ROIs with pain threshold as shown in Fig. 2. The strongest associations include the right insula, right caudate nucleus, bilateral subgenual anterior cingulate cortices (ACC), and bilateral amygdalae. If considering the number of associations as the criterion, the right insula shows the most significant result as four SPN-by-ROI associations can be identified. Moreover, the left insula also shows identification with SNP rs16907751 at the p≤10^-6^ significance threshold. Though the other three SNPs cannot pass the given threshold test, the colormap shows that they also have relatively stronger associations with the left insula than other ROIs. From Fig. 3, some structured patterns can be observed. On the one hand, in the imaging domain, the hierarchical clustering result on the x-axis shows the symmetrical property of bilateral brain hemispheres. Some of these ROIs, such as the insula, thalamus, ACC, and amygdala, belong to the conventional pain matrix, which is involved in the modulation of pain from top-down cognition and bottom-up tactile stimuli (Lu et al. 2016). Meanwhile, the selection of four SNPs (rs10504722, rs16907751, rs396760, and rs187974) all associate with both left and right insula cortexes. It is possible to identify underlying brain connectivity between bilateral regions related to genetic variation. On the other hand, in the genomic domain, these SNPs are hierarchically clustered together to the 8^th^ chromosome as shown in the dendrogram on the y-axis in Fig. 3. This is possibly due to the LD relationships for those SNPs. Many studies about insula and pain have been conducted in recent years. For instance, a former VBM-based study found that the morphological changes of the right anterior insular cortex can affect the pain areas of patients with fibromyalgia (Liu et al. 2022). In (Ferraro et al. 2021), the authors implied that the dysregulation of anterior insula activity can be regarded as a possible functional biomarker of chronic pain from a meta-analysis. Based on prior research, our result is reasonable to identify multiple candidate SNPs associated with the insula.

Based on the heatmap and dendrogram results, we can further analyze potential genetic variation and morphological alterations. Since the right insula has the most SNP associations, we select this ROI as the target phenotype to examine its whole genome mapping in Fig. 4. Firstly, the four identified SNPs can be confirmed to be top markers that pass the p≤10^-6^ significance threshold. Secondly, there are several other SNPs from the same chromosome that also show a relatively stronger association with the right insula. This is may also because of the existing LD relationship but cannot be identified from the heatmap. Thirdly, though the given significance threshold cannot be reached, several SNPs from chromosome 3 also show relatively stronger associations with the right insula. It is still worth conducting further validation research on these results.

To verify the identification of the right insula from our result, a correlation analysis was used to reveal a relationship between its GMD and the pain threshold. As shown in Fig. 4(C), one can see an increasing trend in the normalized pain threshold that has an increased density of the right insula. It implies a negative correlation between GMD and pain sensitivity, i.e. one with higher GMD of the insula may less sensitive to pain stimuli. Despite the arguable heterogeneity (Ruscheweyh et al. 2018), it is now generally accepted that pain perception has an association with gray matter density. For example, several independent studies have suggested that chronic pain patients have a decrease in gray matter in common, and the locations overlapped in the insula, cingulate cortex, and dorsolateral prefrontal cortex (Apkarian et al. 2004; Schmidt-Wilcke et al. 2005; Schmidt-Wilcke et al. 2006; Draganski et al. 2006; Kuchinad et al. 2007; Rodriguez-Raecke et al. 2009). Meanwhile, the morphological changes of gray matter can be observed in phantom pain patients due to amputation and spinal cord injury (Draganski et al. 2006; Wrigley et al. 2009). One possible explanation is that cell atrophy and the decrease in cell size or blood volume can cause stronger pain syndromes (May et al. 2008). Although there is no conclusive research, our result implies that the structural alteration of insula GMD may affect individual pain sensitivity.

A refined statistical analysis based on a chosen SNP-by-ROI association, i.e. the right insula and the SNP rs187974 was further performed. Fig. 5 shows the result of a two-way ANOVA, where two genotypes TT and TC together as TX due to the limited number of TT subjects. The first observation of Fig. 5 is that the TX group has a lower density of gray matter than the CC group for both high and low sensitivity subject groups with p=0.0105. This suggests that the carrier of allele T at SNP rs187974 may lead to a lower GMD in the right insula than in CC subjects. Another observation from Fig. 5 is that the GMD in the high pain sensitivity group is slightly lower than the low sensitivity group, with a p-value of less than 0.05. This statistical result verifies the finding in Fig. 4(c) that subjects with lower GMD may be more sensitive to pain. A detailed analysis was further conducted on the subregions of the insula, including the PI, dAI, and vAI. The result in Fig. 6 shows that the SNP rs187974 is only found to be associated with dAI. This could suggest that the GMD of dAI, but not the entire insula, may affect the individual pain perception. In line with our findings, (Wiech et al. 2010) have demonstrated that the anterior insula encodes the perceived threat value of an upcoming laser stimulation before a stimulus encounter. Evidence was provided that the anterior insula structure plays a pivotal role in integrating information about the threat level of stimulation into the decision about pain. Many previous studies have also indicated that the anterior insula can anatomically integrate different types of information into sensory processing (Mesulam and Mufson 1982; Paulus and Stein 2006). Together with our results, these findings may imply that the anterior insula is a key role in individual interoception that involves sensation monitoring. Further beyond the imaging dimension, our study also shows other possible bases for such findings from the genetic dimension. From Fig. 6, statistical results show a significant association between the other three SNPs and dAI. The two-way ANOVA reveals that the GMD of dAI has statistically significant effects on both genotypes and pain sensitivity. However, there is not a statistically significant interaction between genotype and pain sensitivity from all these results. The SNPs rs10504722 and rs187974 belong to the gene ZBTB10 (Zinc Finger and BTB Domain Containing 10). The genes of the other two SNPs rs16907751 and rs396760 are limited known, it is most likely that they also belong to this same gene due to the hierarchical clustering result from Fig. 3 and the LD relationship. Though indeed a literature search did not reveal any knowledge associating ZBTB10 and pain sensitivity, additional analyses on such SNPs and genes appear possible for future study.

### 4.3 GWAS of functional brain phenotypes

As we can see from Fig. 7, two SNPs rs2267447 and rs1081003 can be identified to be associated with the rs-FC between the right insula and left putamen. The main effect analysis of two-way ANOVA reveals that both genotype and individual pain sensitivity have a statistically significant effect on rs-FC strength. More specifically, the carrier of the C allele of rs2267447 and A allele of rs1081003 may lead to a lower rs-FC value. Meanwhile, the rs-FC in the high pain sensitivity group shows a lower value than in the low sensitivity group. The result implies that the dysregulation of rs-FC between the right insula and right putamen may perform as a functional marker for pain sensitivity. On the other hand, the identified two SNPs rs2267447 and rs1081003 both belong to the gene CYP2D6 (Cytochrome P450 Family 2 Subfamily D Member 6). It encodes the enzyme that is highly expressed in areas of the central nervous system. The cytochrome P450 family proteins are monooxygenases that catalyze many reactions involved in drug metabolism and synthesis (Bertilsson et al. 2002). The CYP2D6 gene is known to be highly polymorphic and is critical for opioid therapy to improve pain management in clinical scenarios (Ruano et al. 2018; Smith et al. 2019). It remains unknown how this gene could change the rs-FC with the insula. However, in the present study, such an identification suggests that the CYP2D6 gene could be a potential therapeutic target for pain modulation.

### 4.4 Limitations and future directions

Though the scale of our self-collected dataset can be considered as large in the field of pain sensitivity research, the number of included subjects is still relatively low for a brain-wide-genomewide study. This first limitation does not allow us a further validation study to divide the subjects into a test group and a validation group. Consequently, in the heatmap result, the significance threshold level at p≤10^-6^ was reported. Unlike the conventional p≤10^-7^ level, a less stringent threshold was used to identify more potential SNP and imaging QT associations. Though a large dataset can usually lead to more robust results, our findings can also be supported by many other studies on the topic of pain. Future studies could incorporate additional participants in the GWAS research to examine the effects of SNP and target imaging phenotypes. Secondly, this study did not consider the gene expression of the identified SNPs nor the gene-gene interactions. These are very important topics for understanding the pathway of pain modulation and the mechanisms of the difference in individual pain sensitivity. Future analysis should employ the gene expression dataset such as AHBA for more comprehensive research to reveal more important factors in pain sensitivity. Thirdly, the AAL3 atlas was used in this study to create ROIs. In the future, a more detailed atlas or probabilistic atlas can be employed to enhance the resolution of imaging phenotype.

## 5 Conclusion

To conclude, we carried out a whole-genome-whole-brain pain sensitivity study in a self-collected dataset with 432 normal participants. We provide evidence that the right insula and its subregion dAI are significantly associated with individual laser pain sensitivity and can play important role in pain perception. Moreover, GWAS and a series of refined analyses demonstrated potential key loci and their associated brain structural and functional phenotypes. Some of the encouraging results and findings are in line well with the existing literature. It indicates that the existence of genetic variants may contribute to individual differences in pain sensitivity. This study is one of the few to investigate the brain structural, functional, and genetic basis of pain sensitivity using an imaging genetic approach. Overall, our results provided several potential quantitative trait loci associated with structural and functional phenotypes of pain sensitivity. These findings may provide more insight into the mechanisms of underlying pain sensitivity in humans, and may also inform the future development of clinical precision pain modulation.

